# Genome-scale mutational signatures of aflatoxin in cells, mice and human tumors

**DOI:** 10.1101/130179

**Authors:** Mi Ni Huang, Willie Yu, Wei Wei Teoh, Maude Ardin, Apinya Jusakul, Alvin Ng, Arnoud Boot, Behnoush Abedi-Ardekani, Stephanie Villar, Swe Swe Myint, Rashidah Othman, Song Ling Poon, Adriana Heguy, Magali Olivier, Monica Hollstein, Patrick Tan, Bin Tean Teh, Kanaga Sabapathy, Jiri Zavadil, Steven G. Rozen

## Abstract

Aflatoxin B1 (AFB1) is a mutagen and IARC Group 1 carcinogen that causes hepatocellular carcinoma (HCC). Here we present the first whole genome data on the mutational signatures of AFB1 exposure from a total of > 40,000 mutations in four experimental systems: two different human cell lines, and in liver tumors in wild-type mice and in mice that carried a hepatitis B surface antigen transgene – this to model the multiplicative effects of aflatoxin exposure and hepatitis B in causing HCC. AFB1 mutational signatures from all four experimental systems were remarkably similar. We integrated the experimental mutational signatures with data from newly-sequenced HCCs from Qidong County, China, a region of well-studied aflatoxin exposure. This indicated that COSMIC mutational signature 24, previously hypothesized to stem from aflatoxin exposure, indeed likely represents AFB1 exposure, possibly combined with other exposures. Among published somatic mutation data, we found evidence of AFB1 exposure in 0.7% of HCCs treated in North America, 1% of HCCs from Japan, but 16% of HCCs from Hong Kong. Thus, aflatoxin exposure apparently remains a substantial public health issue in some areas. This aspect of our study exemplifies the promise of future widespread resequencing of tumor genomes in providing new insights into the contribution of mutagenic exposures to cancer incidence.

## Introduction

Many mutagens and mutagenic processes imprint a characteristic "extended mutational signature" that comprises single nucleotide substitutions in the context of the preceding and following bases, as well as additional features, including transcription and replication strand bias, association with indels or dinucleotide substitutions, and so forth. For example, ultraviolet light tends to induce TCC > TTC mutations across the entire genome, and aristolochic acid induces CAG > CTG mutations (Alexandrov et al. 2013; Hoang et al. 2013; Poon et al. 2013). Both show transcriptional strand bias but negligible replication strand bias, and neither is associated with elevated levels of indel mutations. Unlike aristolochic acid, however, UV light induces high levels of CC > TT dinucleotide substitutions. Elucidation of mutational signatures has been driven by computational analyses of somatic mutations in thousands of tumors, followed by integration with clinical information and prior knowledge of mutational mechanisms (Alexandrov et al. 2013). There are now 30 widely accepted signatures, with a variety of known, suspected, or unknown causes (Wellcome Trust Sanger Institute 2016). These signatures hold promise in molecular cancer epidemiology because they bear witness to mutagenic exposures and because they illuminate endogenous mutagenic processes and mechanisms of DNA damage and repair (Alexandrov and Stratton 2014; Helleday et al. 2014; Poon et al. 2014; Nik-Zainal et al. 2015; Hollstein et al. 2016).

Despite the progress in computational analysis of mutational signatures, the experimental delineation of extended mutational signatures across whole exomes or whole genomes is essential because the causes of many computationally discerned signatures are unknown and because it is possible that several mutagenic processes or exposures could generate similar signatures. While exome- or genome-wide extended mutational signatures from several metazoan experimental systems have been reported (Meier et al. 2014; Olivier et al. 2014; Severson et al. 2014; Nik-Zainal et al. 2015; Poon et al. 2015; Blokzijl et al. 2016; Zamborszky et al. 2016), development of systems that robustly recapitulate *in-vivo* human mutagenesis remains a challenge.

To study the connections between experimentally and computationally derived signatures, we focussed on the mutagenic carcinogen aflatoxin B1 (AFB1). AFB1 is considered the most important of several co-occurring aflatoxins that are produced by molds growing in grain, peanuts, or other food. Aflatoxin exposure is thought to be a major public health risk in parts of Africa and Asia, partly by synergizing with hepatitis B to dramatically increase risk of hepatocellular carcinoma (HCC) (Kensler et al. 2003; IARC Working Group on the Evaluation of Carcinogenic Risks to Humans 2012). However, to our knowledge, the extended mutational signatures of aflatoxins have been little studied in the context of normal metazoan DNA repair: a study in *Caenorhabditis elegans* examined the AFB1 mutation signature based on 31 single base-substitution mutations (Meier et al. 2014), and ultra-deep sequencing of 6.4 kilobases in AFB1 exposed mice inferred an AFB1 signature from 397 mutations (Chawanthayatham et al. 2017).

Conversely, a computationally extracted signature of aflatoxin exposure in HCCs, termed COSMIC Signature 24, has been described based on computational analysis of somatic mutations from the whole-exome sequencing (WES) of a 11 tumors for which aflatoxin exposure was inferred from African or Asian origins or “race” and, in some HCCs, the presence of the *TP53* R249S mutation, which occurs in roughly half of aflatoxin exposed HCCs (Hollstein et al. 1991; Montesano et al. 1997; Ming et al. 2002; Sun et al. 2011; Schulze et al. 2015). In addition, while the current paper was in review, a publication on HCCs from Qidong County, a region of known aflatoxin exposure, extracted a signature similar to COSMIC Signature 24 from HCC somatic mutations (Zhang et al. 2017).

Here we report a first-of-its-kind multi-system strategy for the experimental characterization of the extended mutational signature of AFB1 in human cell lines and in mouse models of AFB1-induced HCC. We further examine the relationships between experimentally determined signatures and mutational signatures in multiple human HCCs with likely dominant aflatoxin exposure and provide a broader overview of the prevalence of likely aflatoxin exposure and provide a broader overview of the prevalence of likely aflatoxin exposure in published whole-genome sequence (WGS) data from human HCCs (Figure 1).

**Figure 1.**
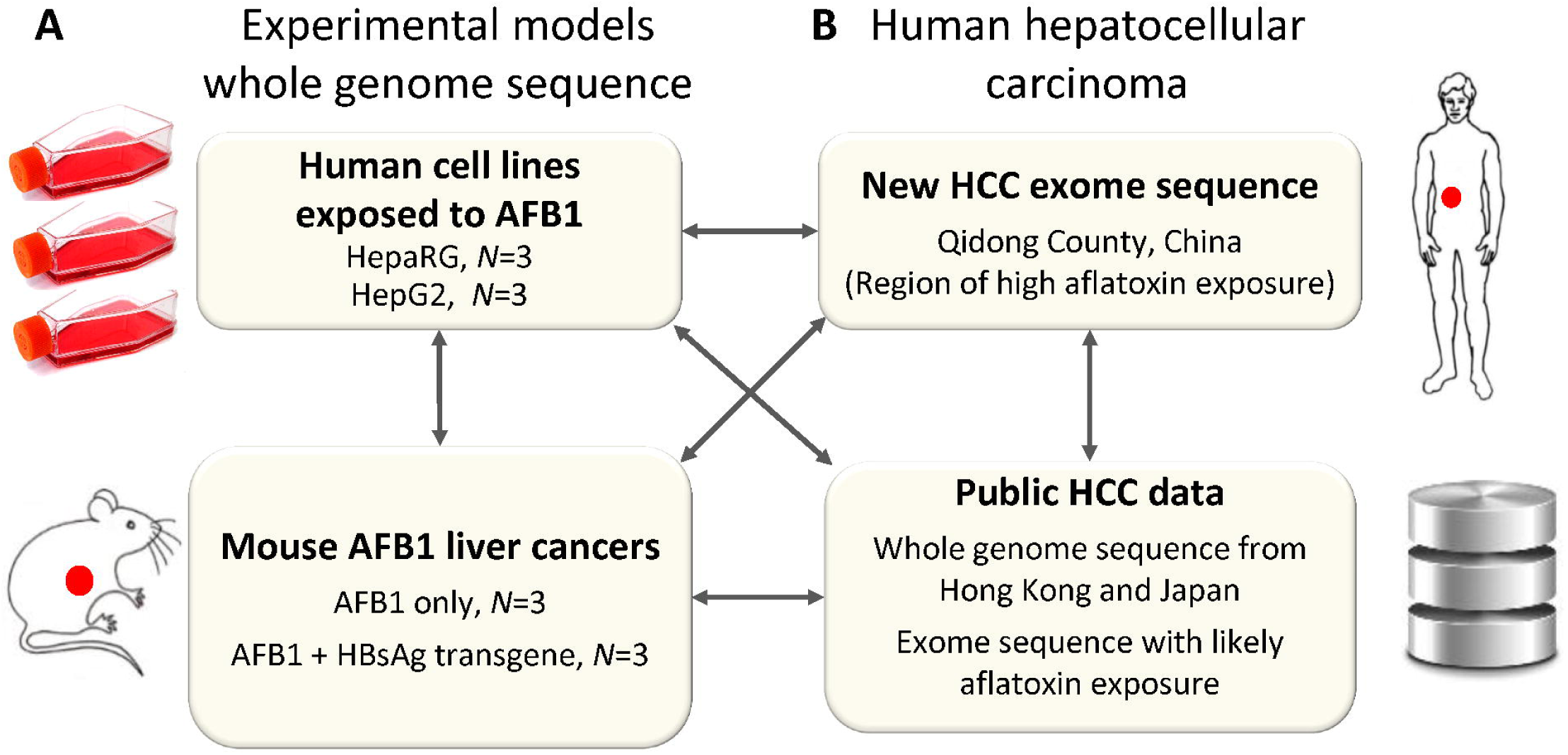
Study design. **(A)** We experimentally elucidated the mutational signature of AFB1 based on whole genome sequence in four experimental systems: two different AFB1-exposed human cell lines, liver tumors in AFB1-exposed wild-type mice and in AFB1-exposed mice that carried a hepatitis B surface antigen transgene. In total we examined 48,000 mutations in experimental systems including mutations in cell lines and somatic mutations in mouse tumors. **(B)** We integrated these experimental results with newly generated genomic HCC data from a geographical region of known aflatoxin exposure and with additional, publicly available human HCC data.

## Results

### Mutation spectra of AFB_1_ exposed human cell lines

To investigate the extended mutational signature of AFB1, we first whole-genome sequenced two AFB1-exposed HCC cell lines, HepaRG and HepG2 (Figures 1, 2). Mutation patterns were remarkably stable across 3 replicates within each cell line (average pairwise Pearson correlation > 0.97, Supplementary table 1, Supplementary figure 1). Mutation patterns were also similar between HepG2 and HepaRG, but slightly less than within each cell line (average pairwise Pearson correlation 0.94).

G > T mutations predominated, accounting for 68% and 50% of the total G > N mutations for HepaRG and HepG2 respectively (Supplementary table 2). G > A mutations were the next most abundant, followed by G > C mutations. The predominance of G > N mutations and the relative proportions of G > T, G > A, and G > C mutations were broadly consistent with previous experimental studies (Foster et al. 1983; Bailey et al. 1996). In the extended mutational signatures, TGC > TTC mutations were most frequent among the G > T mutations, and AGC > ATC and TGG > TAG mutations were also noticeably enriched (Figure 2, Supplementary figure 1). There was strong enrichment of G > T mutations on the non-transcribed strands of genes (Figure 2B, Supplementary figure 2). The primary mutagenic mechanism of AFB1 is formation of adducts on guanines (Guengerich et al. 1998), and the relative paucity of mutations on the non-transcribed strands is presumably due to transcription coupled nucleotide-excision repair (TC-NER) of damage occurring on guanines (Fousteri and Mullenders 2008). Furthermore, consistent with TC-NER activity, the level of bias was correlated with mRNA levels in the respective cell lines: in HepaRG only 14% of the mutations in the most highly expressed genes were on the transcribed strand versus 47% in the least expressed genes (Supplementary table 3). In HepG2, the corresponding figures were 11% and 53%. There is also prior evidence that the effectiveness of TC-NER decreases from the 5' to 3' ends of transcripts (Conaway and Conaway 1999; Hu et al. 2015). Consistent with this, and further supporting the involvement of TC-NER in the repair of AFB1-guanine adducts, the ratio of non-transcribed to transcribed-strand G > T mutations decreased from 2.6:1 in the first 100 kb of transcripts to approximately 1.5:1 in the 200 kb centered on 0.5 Mb (Supplementary figure 3, p < 7×10^-8^ for HepaRG and p < 1.6×10^-4^ for HepG2, by logistic regression).

**Figure 2.**
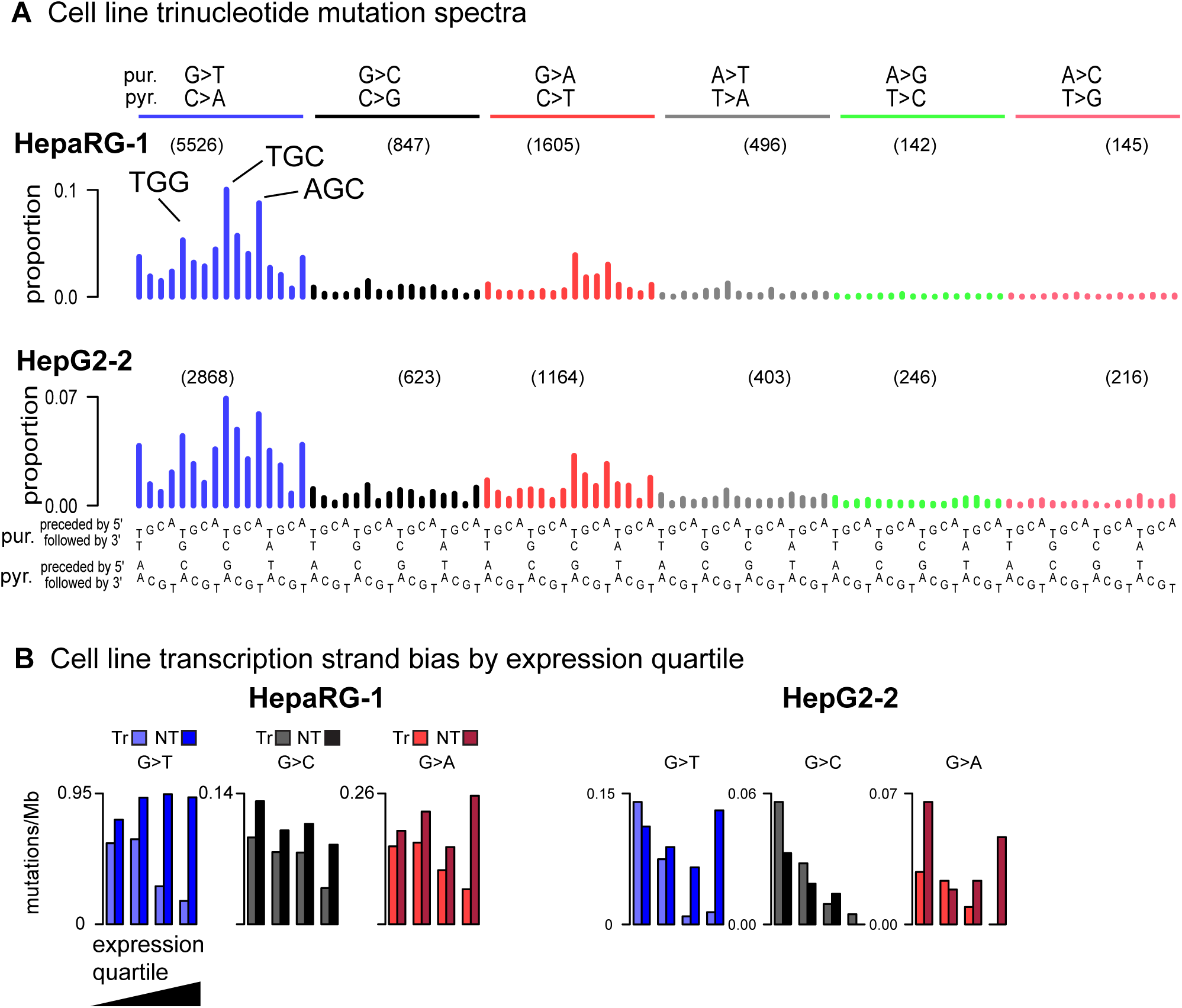
**(A)** Representative human-cell line trinucleotide mutation spectra grouped by mutations from guanine (G > T, G > C, G > A) and adenine (e.g. A > T, A > G, A > C). The most frequent G > T mutations are indicated (TGG > TTG, TGC > TTC, AGC > ATC). The number of mutations in in each mutation class (e.g. G > T) are indicated in parentheses. As there was little variation between replicates within each cell line, we show all individual spectra in Supplementary figure 1. **(B)** Extreme transcription strand bias for genes with high expression levels; see Supplementary figure 2 for transcription strand bias for all cell line replicates.

We also looked for variation in mutation intensity across the genome as a function of replication time or replication strand. In HepG2 but not HepaRG there was significant enrichment for mutations in late-replicating regions (> 56% of G > T mutations in late regions, p < 6×10^-6^ in any HepG2 replicate, Supplementary table 4.) We are unable to explain lack of replication-time bias in the HepaRG findings. However, in both cell lines, we observed slight but consistent enrichment of G > T mutations on the leading strand (52.2%, p < 0.0004 for HepaRG and 51.6%, p < 0.032 for HepG2, Supplementary table 5).

We also analyzed the AFB1 mutation data for dinucleotide mutations from guanines, that is, substitutions of two nucleotides: GN > NN or NG > NN. These "G dinucleotide mutations" are associated with COSMIC Signature 4, which is associated with tobacco smoking and also has a high proportion of G > T mutations (Alexandrov et al. 2013). In the AFB1-exposed cells, the ratios of the number of G-dinucleotide mutations to G > N mutations were < 0.011 (Supplementary table 2), lower than any of the HCCs with COSMIC Signature 4 in (Fujimoto et al. 2016) (minimum 0.0134).

### Mutation spectra of mouse AFB1 HCC-like liver tumors

We studied mutation spectra in two mouse models of aflatoxin-associated HCCs. One model consisted of tumors from mice exposed only to AFB1. The other model consisted of tumors from mice bearing the weakly oncogenic hepatitis B surface antigen (HBsAg) transgene (Chisari et al. 1985) and exposed to AFB1, a system designed to model the synergistic carcinogenic effects of hepatitis infection and aflatoxin exposure (Kensler et al. 2003; IARC Working Group on the Evaluation of Carcinogenic Risks to Humans 2012). The WGS-based mutation spectra of all the mouse tumors, like the human cell lines, were dominated by G > T mutations and also showed noticeably high numbers of G > C and G > A mutations (Figure 3A,B). However, AFB1+HBsAg tumors had far fewer G > N mutations (mean 468) than the AFB1-only tumors (mean 3,242).

**Figure 3.**
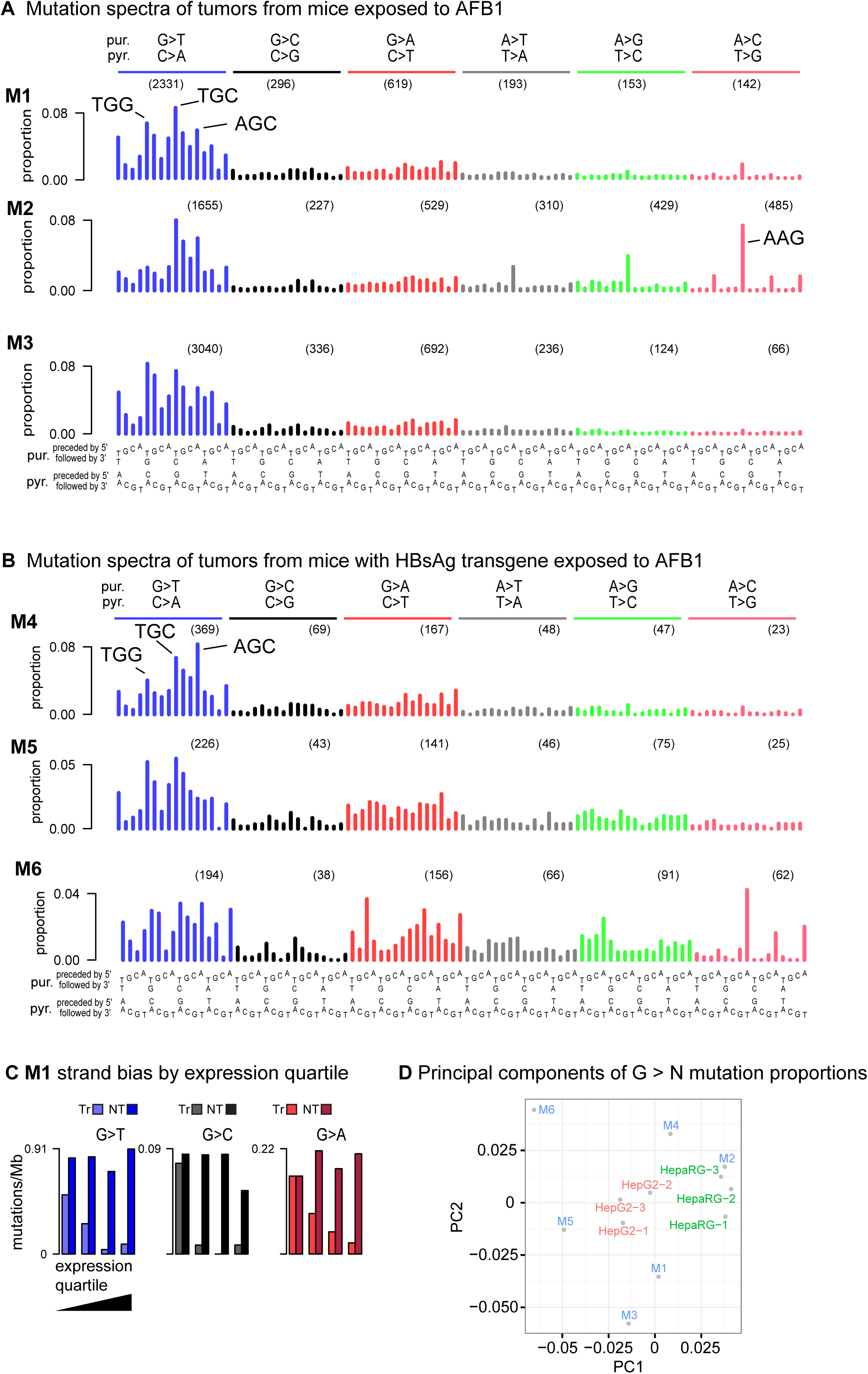
Somatic mutation spectra from HCC-like liver tumors from (**A**) 3 AFB1-exposed mice and **(B)** 3 AFB1-exposed mice with an HBsAg transgene. The latter have only one-tenth as many mutations as tumors from the AFB1-only mice. Spectra in panels A and B were normalized to the trinucleotide frequencies in the human genome. (**C**) Extreme transcription strand bias for all G > N mutations in highly expressed genes in mouse M1. See Supplementary figure 3 for other mice. Because of low mutation count, transcription strand bias is evident only in the G > T mutations in the tumors from the AFB1+HBsAg mice. **(D)** Principal components analysis (PCA) on G > N mutations in trinucleotide context. Replicates of each of the cell lines cluster together, while the mouse tumors are more dispersed in principal components space. The greater dispersion among the HBsAg tumors (M4, M5, M6) is likely due to higher stochastic variance because of much lower mutation counts combined with greater relative contributions from other mutational processes that arose during tumor development.

We observed more variability between spectra among the mouse tumors than among each cell line, with an average Pearson correlation of 0.89 among the AFB1-only tumors compared to 0.97 within each cell line and 0.94 among all cell lines (Supplementary table 1). This was also apparent in a principal components analysis (PCA) (Figure 3D). The AFB1+HBsAg tumors had even more diversity, probably due primarily to the greater relative sampling variation implicit in fewer mutations. Additional variability among the mouse tumors may have been due to additional mutagenic processes before and during cancer development. Notably, in addition to the G > N mutations attributable to AFB1, the spectra from M1, M2, and M6 contained COSMIC Signature 17. This signature, which is of unknown etiology, is characterized by isolated peaks at AAN > ACN, especially at AAG > ACG, and at AAG > AGG (Alexandrov et al. 2013). We considered the possibility that the emergence of COSMIC Signature 17 in these tumors might stem from some somatic mutation affecting DNA repair or activating some endogenous mutagenic process. However, we were not able to identify any somatic mutations in the affected tumors that might account for this additional signature.

To further investigate sources of diversity among the mutation spectra, we stratified the spectra by variant allele frequency (VAF), as variants with lower VAF are often sub-clonal and, as such, are likely to have arisen later in tumor development. We also analyzed the tumors for aneuploidy and chromosomal copy number to allow better inference of clonality from the VAFs (Supplementary figures 4 through 15). AFB1 was administered in a single dose at postnatal day 7, leading to the expectation that all or most AFB1-associated mutations would have occurred prior to tumor initiation and be clonal. Consistent with this, the analysis of the VAFs of G > T mutations indicates that almost all are clonal. Interestingly, the COSMIC Signature 17 mutations were subclonal and therefore presumably occurred later in tumor development. There was also a diffuse signature similar to COSMIC Signature 5 noticeable in mutations with VAFs < 0.2 in M3 (Supplementary figure 8).

Despite the greater diversity in mouse tumor G > N spectra, they broadly resembled the cell lines. As in the cell lines, TGC > TTC, AGC > ATC, and TGG > TAG mutations were prominent in all AFB1-only tumors and in the much less mutated AFB1+HBsAg tumors. In the AFB1-only tumors, G > T mutations constituted 72% of the total G > N mutations on average (Supplementary table 2), which was similar to the proportion in the HepaRG cells. In the AFB1+HBsAg tumors, G > T mutations constituted 55% of all G > N mutations, possibly because of G > C and G > A mutations due to factors other than AFB1 exposure. As in the cell lines, the ratio of G dinucleotide mutations to the total number of G > N mutations was low (< 0.0123 in all tumors, Supplementary table 2).

Also like the human cell lines, and consistent with the operation of TC-NER on guanine adducts, the mouse tumors were strongly enriched for G > T mutations on the non-transcribed strands of genes, and the level of enrichment was higher in more highly expressed genes: among the AFB1-only tumors, an average of only 7% of G > T mutations occurred on the transcribed strands of genes in the top expression quartile, and this pattern of expression-level-correlated strand bias extended to G > C and G > A mutations (Figure 3C, Supplementary figure 16, Supplementary table 3). Among the AFB1+HBsAg tumors, G > T strand bias in the top expression quartile was also detectable in the aggregate, with 26% of these mutations on the transcribed strand. We hypothesize that the weaker strand bias reflects the presence of other mutational processes that also generate G:C > T:A mutations, which would have had more relative impact in the low-mutation AFB1+HBsAg tumors. Also like the human cell lines, and consistent with the operation of TC-NER of guanine adducts, transcription strand bias decreased with distance from the 5' end of the gene: the ratio of non-transcribed to transcribed-strand G > T mutations decreased from 4.1 in the first 100 kb to 1.5 at the 200 kb centered at 0.5 Mb (p = < 1.7×10^-8^ by logistic regression, Supplementary figure 17).

We were unable to assess whether there was a difference in mutation intensity between early and late replicating regions or between leading and lagging replication strands, as we could not locate replication timing data for mouse hepatocytes.

### Mutation spectra of likely aflatoxin-associated human HCCs

We integrated the experimental results from cell lines and mice with somatic-mutation spectra from newly sequenced HCCs from Qidong, China, a region of well-studied high aflatoxin exposure (Szymanska et al. 2009; Kensler et al. 2011). The HCCs from Qidong were also selected for somatic *TP53* R249S mutations, which is considered indicative of AFB1 exposure (Hollstein et al. 1991; Montesano et al. 1997; Sun et al. 2011). The Qidong HCCs had high proportions of G > T mutations and transcription strand bias skewed toward highly expressed genes, consistent with aflatoxin exposure (Supplementary figures 18, 19, Supplementary table 3). In this exome data, there were too few mutations to assess whether transcription-strand bias decreased along the length of transcripts (Supplementary figure 20).

Of note, one HCC from Qidong (PT4, Supplementary figure 18) had a prominent A > T mutation pattern indicative of exposure to aristolochic acid (Hoang et al. 2013; Poon et al. 2013). However, considered apart from A > T mutations, the G > N pattern in this HCC was consistent with aflatoxin exposure, suggesting past co-exposure to these two strong carcinogens.

We next analyzed additional, publicly available HCC spectra, some with previously hypothesized aflatoxin exposure, from the following sources: (1) 11 HCCs (1 WGS, 10 WES) with African or Asian origins or "race" that had patterns of mutations thought to be due to aflatoxin exposure (Schulze et al. 2015), (2) WGS of 79 hepatitis-B positive HCCs from Hong Kong (Sung et al. 2012) and (3) WGS of 268 HCCs from Japan (Fujimoto et al. 2016). PCA of the proportions of G > T mutations in trinucleotide context separated the experimental spectra from the vast majority of the HCC spectra by lower values of principal component 1 (PC1, Figure 4). To select likely aflatoxin-affected HCCs for further study, we used as a threshold the maximum value of PC1 across the experimental data (Figure 4, M3's PC1 value, indicated by a vertical green line). We selected the 8 WGS HCCs that exceeded this threshold for initial study. Among these 8 was DO23048, which was identified in (Schulze et al. 2015) from WES data as probably aflatoxin-exposed. All 6 of the WES HCCs from Qidong exceeded this threshold. Furthermore, the 10 presumed aflatoxin-related WES tumors in (Schulze et al. 2015) also exceeded this threshold and showed patterns of G > T and G > N mutations and transcriptional strand bias similar to that observed in the Qidong HCCs (Supplementary figures 21 through 24). An additional PCA, based on the proportions of G > N mutations in trinucleotide context, likewise placed the 8 selected aflatoxin-linked WGS HCCs and the WES HCCs from Qidong or identified in (Schulze et al. 2015) near the experimental samples and away from the vast majority of the remaining HCCs (Supplementary figure 25). To gain insight into possible aflatoxin exposure in North America, we repeated the PCA after adding the full set of HCC WES somatic mutations from The Cancer Genome Atlas (TCGA) (The Cancer Genome Atlas 2017). This flagged 5 additional possible aflatoxin-exposed HCCs. However, examination of the spectra suggested that they did not reflect detectable aflatoxin exposure, because of minimal transcriptional strand bias and because of the patterns of G >T mutations in trinucleotide context (Supplementary figure 26). Thus, as far as can be determined from the TCGA WES somatic mutations, the rate of aflatoxin exposure in HCCs treated in North American HCCs was 2 in 289, or 0.7%.

**Figure 4.**
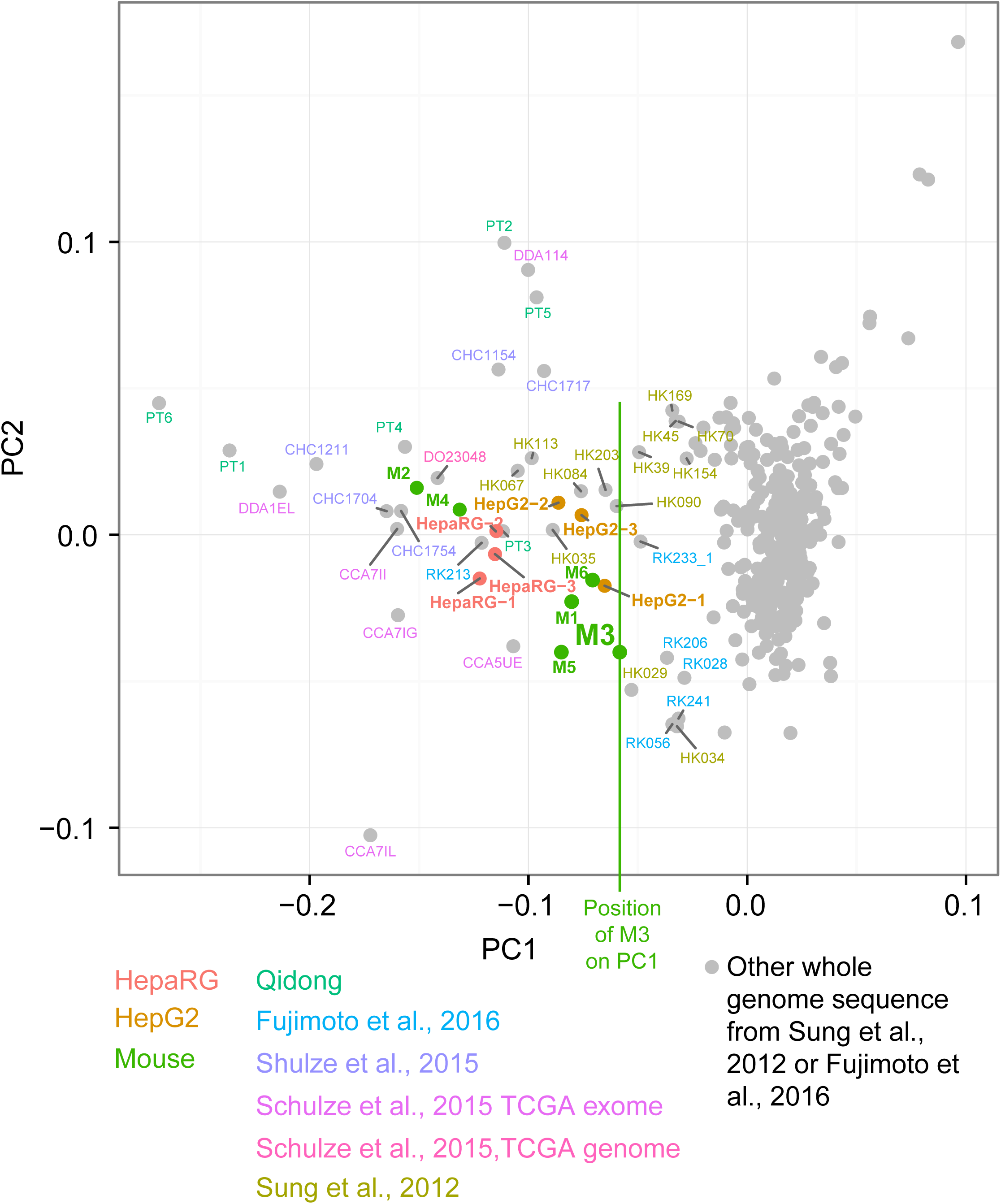
Selection of likely aflatoxin-associated HCCs by principal components analysis on the proportions of G > T mutations in trinucleotide context. Vertical green line indicates the value of M3 in PC1, which was used as a threshold for selecting WGS HCCs likely exposed to aflatoxins for further study: One HCC identified from WES data in (Schulze et al. 2015) for which WGS data was subsequently available (DO23048), 6 from (Sung et al. 2012), and one HCC from (Fujimoto et al. 2016). As expected due to the higher relative sampling variance in WES spectra due to small numbers of mutations, these were more variable than WGS spectra.

As in the experimental systems, the spectra of the 8 WGS HCCs initially selected for analysis had abundant G > T mutations, and consistent with the operation of TC-NER, marked transcription strand bias skewed toward highly expressed genes (Figure 5A, Supplementary figures 27, 28, Supplementary table 3). As was the case for the cell lines and mouse tumors, and again consistent with the operation of TC-NER, transcription strand bias decreased with distance from the 5' end of the gene: the ratio of non-transcribed to transcribed-strand G > T mutations decreased from 2.3:1 in the first 100 kb to 1.3:1 at the 200 kb centred at 0.5 Mb (p = < 3×10^-10^ by logistic regression, Supplementary figure 29). In addition, like the HepG2 cells, all 8 WGS HCCs showed an excess of likely AFB1-associated mutations in late replicating regions of the genome (> 60% in late rather than early replicating regions, p < 10^-12^ for G > T mutations, Supplementary table 4). Also, like the cell lines, the 8 WGS HCCs had slight but consistent excesses of G > T mutations on leading replication strands (Supplementary table 5, 52% of G > T mutations on the leading strand, p = 1.6×10^-7^ by binomial test on aggregated mutation counts).

**Figure 5.**
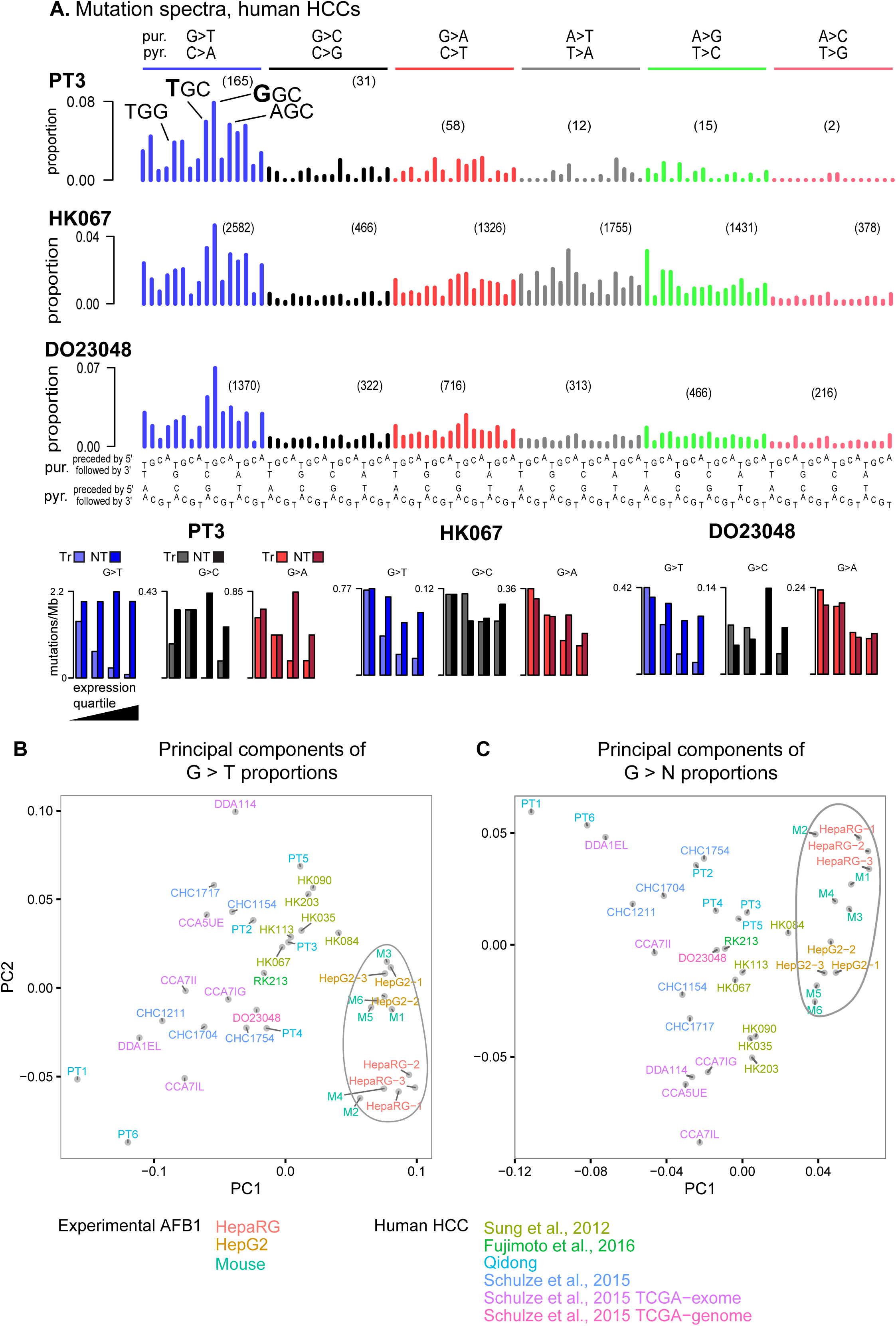
(**A**) Somatic mutation spectra and transcriptional strand bias for initial study of aflatoxin signatures in example human HCCs with likely aflatoxin exposure. (**B**) G > T and (**C**) G > N mutations for AFB experimental data (enclosed in irregular ovals) and HCCs with likely high aflatoxin exposure.

However, the G > T mutations in the HCCs differed from the experimental systems in that the most frequent mutation was GGC > GTC rather than TGC > TTC (Figures 1A, 2A,B). Consistent with this, PCAs that included only the experimental systems and the likely aflatoxin exposed HCCs separated these two groups of samples (Figure 5B,C). We note that the G > N mutation spectra of the 8 selected WGS HCCs were more similar to each other as a group than were the AFB1-only mouse tumors (average pairwise Pearson correlation among the WGS HCCs was 0.92, among the AFB1-only mouse tumors, 0.89). Thus, the G > N mutation spectra of these WGS HCCs may reflect a reasonably uniform underlying set of mutational processes. Among the 8 selected HCCs, G > T mutations accounted for an average of 58% of G > N mutations (Supplementary table 2), somewhat less than for cell line and AFB1-only mouse tumors. The relatively higher proportions of non G > T mutations probably reflects the presence of mutations from other processes, especially among the G:C > A:T mutations, which can be generated by several other mutational processes that operate in HCCs (Wellcome Trust Sanger Institute 2016). Indeed, inspection of the spectra suggests presence of COSMIC Signatures 12 and 16, which are both common in HCC and generate G:C > A:T mutations.

### Decomposition of aflatoxin signatures and prevalence of aflatoxin exposure in HCC

In view of the differences in the G > T spectra between the aflatoxin associated HCCs and all three experimental systems – namely the predominance of GGC > GTC in HCCs rather than TGC > TTC (Figures 1A,2B,C,4A), we used a non-negative matrix factorization techniques to factor the spectra of the AFB1-exposed cell lines and mouse tumors and likely-aflatoxin-exposed WGS HCCs into two signatures based on the 48 G > N mutation classes; we call these two signatures AFB1sig_G>N_ and AFsig2_G>N_ (Figure 6A,B,C). Reconstruction of the observed G>N spectra was quite accurate (Supplementary table 6). Given that COSMIC Signatures 5 or 6 and have been observed in human HCCs we sought to determine if they were affecting signature decomposition. To this end we performed two separate factorizations into 3 signatures. One factorization specified inclusion of COSMIC Signature 5 and the other specified inclusion of COSMIC 6. Neither signature affected AFB1sig_G>N_, but AFsig2_G>N_ was slightly affected (Supplementary table 7). It is possible that AFsig2_G>N_ represents simply a subtraction of AFB1sig_G>N_ from the G > N spectra of HCCs rather than an independent mutational process. Clarification of this will depend on future investigation into the causes of differences between the experimental AFB1 spectra and the spectra in HCCs.

**Figure 6.**
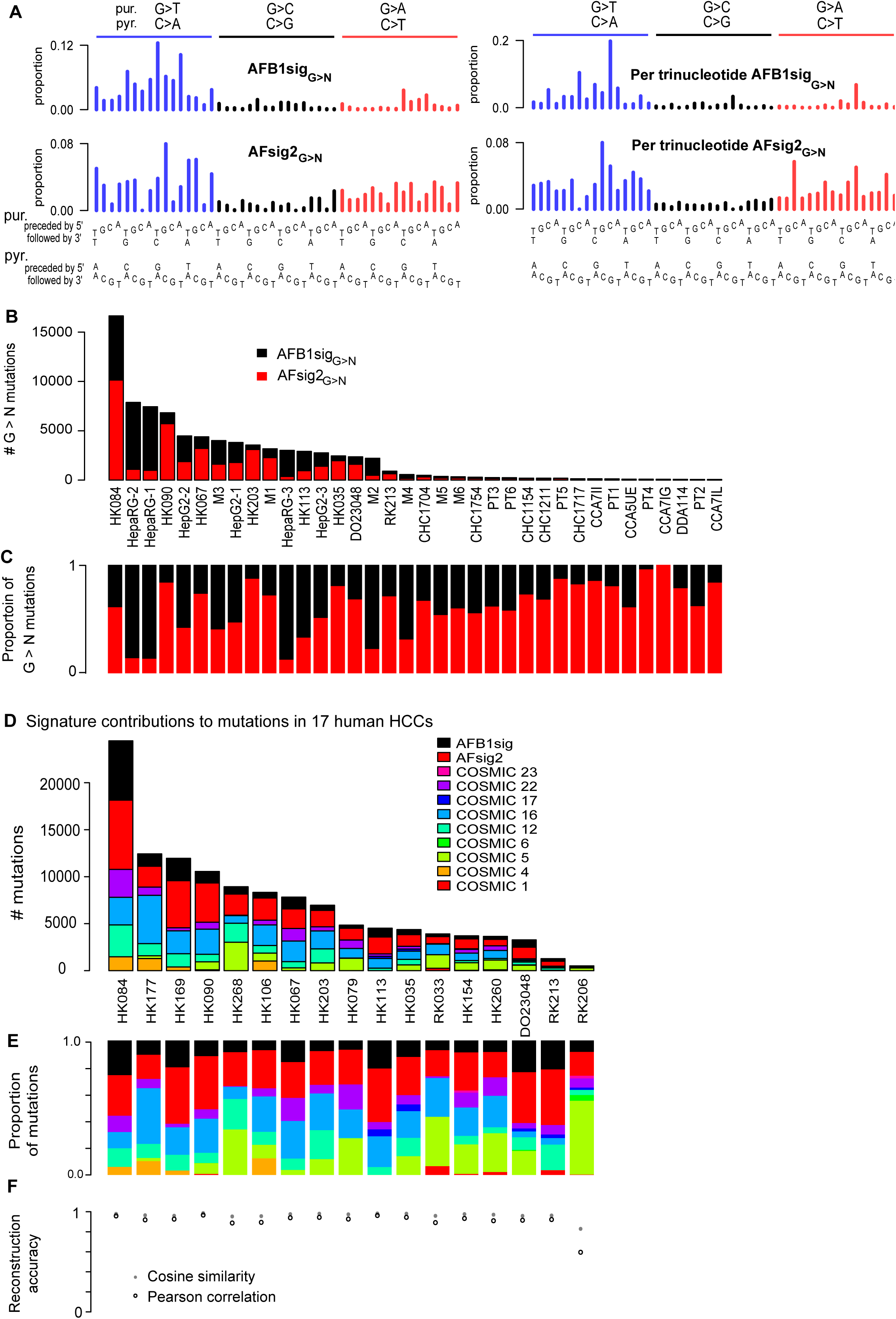
Aflatoxin signatures in human HCCs. (**A**) NMF decomposes the G > N spectra from experimental AFB1 exposure and likely aflatoxin-associated HCCs into two components, denoted AFB1sig_G>N_ and AFsig2_G>N_. The left hand of the panel shows the signatures, as is conventional, based on trinucleotide frequencies in the human genome; the right hand side shows them in frequencies per trinucleotide—equivalent to assuming all trinucleotides are equally common. (**B,C**) AFB1sig_G>N_ (black) almost completely captures the spectra of HepaRG cell lines; AFsig2_G>N_ (red) almost completely captures the spectra of some HCCs (e.g. HK067, HK203), while the mutations in HepG2, the mouse tumors and some HCCs are most accurately reconstructed by a mixture of AFB1sig_G>N_ and AFsig2_G>N_. (**D,E**) Non-negative matrix factorization of selected HCCs using mutational signatures know to occur in HCCs plus extensions of AFB1sig_G>N_ and AFsig2_G>N_ with A > N proportions set to 0, the later denoted AFB1sig and AFsig2. (**D**) Absolute mutation numbers assigned to each signature (**E**) Proportions of mutations assigned to each signature. (**F**) Reconstruction accuracy is generally good with the exception of RK206, which has few mutations and a hard-to-reconstruct mutation spectrum outside of G > T mutations, partly due to spikes at TCA:TGA > TTA:TAA and CTG:CAG > CCG:CGG.

We then used the NMF framework and the experimentally derived AFB1-related signatures to analyze the 79 WGS somatic mutation spectra from (Sung et al. 2012) and the 268 WGS spectra from (Fujimoto et al. 2016) to identify additional HCCs with likely aflatoxin exposure. We analyzed each HCC using COSMIC signatures known to be associated with HCCs, but replacing COSMIC Signature 24 with extensions of AFB1sig_G>N_ and AFsig2_G>N_ to all 96 trinucleotide contexts (with all A > N mutation proportions set to 0). We termed these AFB1sig and AFsig2. We did not include COSMIC Signature 24, because we reasoned that both it and the combination of AFB1sig and AFsig2 captured most aspects of aflatoxin exposure, and we sought to assess the presence of AFB1sig and AFsig2 without interference from the largely overlapping effects of COSMIC 24. Based on this analysis, in addition to the 8 HCCs selected above in Figure 4, nine had substantial evidence for the presence of AFB1sig (i.e. > 5% of the mutations attributed to AFB1sig, Figure 6D,E,F, Supplementary table 8). In total, at least 13 of 79 HCCs from Hong Kong (16%) (Sung et al. 2012) and 3 of the 286 from Japan (1.0%) (Fujimoto et al. 2016) appear to have been exposed to AFB1.

## Discussion

This study combined analysis of experimentally induced AFB1 mutational signatures with computational analysis of mutational signatures in human HCCs thought to have been exposed to aflatoxins. The mutational signatures from all four experimental systems were remarkably similar. NMF and PCA indicate that the 6 *TP53* R249S and hepatitis-B positive HCCs from Qidong were indeed likely exposed to AFB1. NMF and PCA also showed that that HCCs in which COSMIC Signature 24 was previously observed (Schulze et al. 2015) were likely exposed to AFB1. By extension, COSMIC Signature 24 likely represents, at least in part, AFB1 exposure. The evidence for these assertions is as follows. (1) The G > N spectra of the likely-aflatoxin-related HCCs are similar to the experimental spectra in terms of the proportions of G > T, G > A, and G > C mutations. Furthermore, in PCAs on G > T and G > N mutations (Figure 4 and Supplementary figure 13), these HCCs clustered toward the AFB1 experimental systems and away from the overwhelming majority of other HCCs. (2) Both the HCCs and the AFB1 experimental systems showed notable transcription strand bias that strongly skewed toward highly expressed genes and that declined from the 5' to 3' ends of transcripts. (3) The HCCs and the HepG2 cells were enriched for mutations in late replicating regions, and the HCCs and both cell lines were enriched for mutations on the leading replication strand. (We were unable to assess these characteristics in the liver tumors of AFB1-exposed mice.) (4) Both the HCCs and experimental AFB1 systems had low burdens of dinucleotide mutations, lower than any HCC with the smoking-associated COSMIC Signature 4, which might otherwise be difficult to distinguish from aflatoxin exposure. (5) Association of the G > N spectra of the tumors in (Schulze et al. 2015) with aflatoxin was supported with geographical information making exposure plausible. This was also true for the 6 HCCs from Qidong, a region of well-documented aflatoxin exposure. Furthermore, 13 additional HCCs with AFB1sig were from southern China, another region with documented aflatoxin exposure (Wang and Liu 2007). Conversely, only 3 of 268 HCCs from Japan, a region of low exposure, had AFB1sig. (6) Many of the HCCs with strong evidence of aflatoxin exposure also bore the *TP53* R249S mutation, which sometimes indicates aflatoxin exposure in HCCs (Hollstein et al. 1991; Montesano et al. 1997; Sun et al. 2011). The Qidong HCCs were selected based on presence of this mutation, while 6 of the 11 tumors identified in (Schulze et al. 2015) had this mutation, as did 4 of the 12 from (Sung et al. 2012) (Supplementary table 9).

While the prevalence of the TP53 R249S mutation in HCCs with combined hepatitis-B and aflatoxin exposure is biologically remarkable, the sensitivity of this mutation as an indicator of aflatoxin exposure has been difficult to study systematically. Partly this is because peripheral biochemical markers of aflatoxin exposure, such as aflatoxin-albumin or urinary aflatoxin-DNA adducts reflect recent exposure. This makes it difficult to assess the strength of the association between the R249S mutation and long-term aflatoxin exposure (Groopman et al. 1985; Wild et al. 1986; Egner et al. 2006). Estimates of the proportion of tumors with R249S mutations in HCCs with hepatitis-B and aflatoxin-exposure have varied by study, and may depend on the level of aflatoxin exposure (Hollstein et al. 1991; Montesano et al. 1997; Ming et al. 2002; Szymanska et al. 2009; Sun et al. 2011; Chittmittrapap et al. 2013; Kew 2013). Thus a detection method based on robust, well-defined mutational signatures would be of interest for identifying HCCs with aflatoxin exposure but lacking the R249S mutation. The WGS-based approach presented here is a step in that direction, promising specificity if not sensitivity. Given that other exposures, including tobacco smoking and oxidative damage of guanines also affect the pattern of G > T mutations and could obscure evidence of weak AFB1 exposure, further work in this area is needed.

An unexpected finding in the present study was that the AFB1 experimental signatures were different from those in the HCCs, visible mainly in the shift of the most frequent mutations from TGC > TTC in the experimental systems to GGC > GTC in the HCCs (Figures 2, 3, 5). These differences seem unlikely to stem from fundamental biological differences in cellular susceptibility in the experimental systems compared to the HCCs, because the experimental spectra are all relatively similar despite comprising two different human cell lines and *in vivo* tumors from mice both with and without the HBsAg transgene (Figure 5B,C, Supplementary table 1). We also note that the signature of AFB1 mutagenesis inferred from 397 mutations in 6.4 kilobases of sequence in AFB1 exposed mice, (Chawanthayatham et al. 2017) is similar to the AFB1sig_G>N_ reported here (cosine similarity 0.96). (Chawanthayatham and colleagues show mutation frequencies per trinucleotide; for comparison, the right hand side of Figure 6A shows AFB1sig_G>N_ using the same conventions.) One potential reason for differences between experimental AFB1 signatures and the computationally extracted signatures from the human HCCs might be that human exposure does not consist of pure AFB1, but rather a mixture of aflatoxins, including aflatoxins G1, B2, or G2, possibly co-occurring with other mycotoxins. Another potential reason might be differences in dosage, as HCCs might reflect lower, chronic exposures over decades rather than the short periods of high exposure in the experimental systems. A final potential reason might be interaction of aflatoxin with hepatitis; although the mice with the HBsAg transgene had spectra similar to the other experimental AFB1 spectra, this model may not recapitulate all the interactions between AFB1 and hepatitis that contribute to human hepatocellular carcinogenicity.

Beyond the differences between the experimental AFB1 spectra and the HCCs, there was considerable diversity among the mouse tumors and there were subtle but apparently consistent differences between the spectra in HepaRG and HepG2 cells. Diversity among the mouse tumors is partly due to additional mutagenic processes that arose before and during cancer development. This is especially evident in the action of COSMIC Signature 17, probably later in tumor development, in mice M1, M2, and M6 (Supplementary figures 6, 8, 12). In addition, the numbers of mutations in the tumors in the AFB1-exposed mice with the HBsAg transgene were substantially lower than those in the wild-type mice, even though AFB1 exposure and age at tumor sequencing were the same in both groups. Possibly AFB1 was unevenly distributed throughout the liver, leading to varying levels of exposure and mutagenesis among cells. Under this hypothesis, we speculate that in the HBsAg mice, oncogenesis required few AFB1 mutations in addition to the HBsAg transgene, while for wild-type mice, oncogenesis occurred only in the most heavily mutated cells. This scenario would be consistent with the observation of smaller and far fewer tumors in AFB1-treated wild type mice than in AFB1-treated HBsAg mice (Teoh et al. 2015).

Differences between HepaRG and HepG2 may reflect differences in the metabolism of AFB1 between to the two cell lines or differences in intensity of mutagenesis. HepaRG and HepG2 differ in levels of and inducibility of some of the cytochrome P450s that metabolize AFB1 to both the adduct-forming AFB1-exo-8,9-epoxide and to other, less mutagenic metabolites (Hart et al. 2010; Gerets et al. 2012). For example, levels of CYP3A4 and CYP3A5 are much higher in HepaRG than in HepG2. However, the proximal adduct forming compound is thought to be AFB1-exo 8,9-epoxide, regardless of which cytochrome P450s are expressed. Consequently, to the best of the field's understanding, differences in P450 profiles would affect mainly the dosage of AFB1-exo 8,9-epoxide. In addition, HepaRG cultures were treated with higher concentrations of AFB1 than the HepG2 cultures. Thus, dosage effects might account for the subtle differences between HepaRG and HepG2 mutational spectra. However, we cannot exclude the possibility of other, uncharacterized differences in AFB1 mutagenesis between the two cell lines that could account for these differences.

In conclusion, our multi-system approach to linking mutational signature to carcinogen exposure generally confirmed the presence of an aflatoxin signature but also pointed to some of the complexity of aflatoxin and AFB1 carcinogenesis. Our findings further indicate that even for a relatively mutagenic compound such as AFB1, WGS yields more robust views of mutational signatures than WES analysis, because of the higher relative stochastic variability inherent in the smaller numbers of exonic mutations (Supplementary table 10). We propose that the described multi-system approach to the experimental study of carcinogen exposure should be developed further and can be used more broadly to understand the mechanisms of environmental mutagenesis and to provide support for molecular epidemiology studies ultimately aimed at cancer prevention.

## Methods

### Human cell lines

We assessed AFB1 mutations in two human cell lines. HepaRG (Invitrogen) cells were derived from an HCC and can be differentiated to cells that resemble adult hepatocytes; we cultured and differentiated HepaRG cells as described previously (Gripon et al. 2002). HepG2 was derived from a hepatoblastoma (Ostlund et al. 1996); we cultured it in minimum essential medium supplemented with 10% fetal calf serum, nonessential amino acids, 100 units/ml penicillin, and 100 ug/ml streptomycin. We determined the AFB1 (A 6636, Sigma) half maximal inhibitory concentration (IC50) in these cell lines as shown in Supplementary figure 30. To generate clones we treated HepaRG at 30 uM and HepG2 at 20 uM every 3 days for 45 days. We then separately sequenced three clones each of treated HepaRG and HepG2 cells.

### Mouse models

Details of the mice, their treatment, and similarity of the mouse liver tumors to human HCCs are provided in (Teoh et al. 2015). Briefly, we studied tumors from 6 C57BL/6J mice, 3 of which bore an hepatitis B surface antigen (HBsAg) transgene (Chisari et al. 1985). The mice received 1 peritoneal injection of 19 nmol/g AFB1 at day 7 after birth and were sacrificed at 15 months. Animal experiments were approved by and performed in accordance with the guidelines of the SingHealth Animal Care and Use Committee.

### Sources of newly sequenced human tumor and normal pairs

We studied archived paraffin blocks of 6 HCCs and corresponding non-tumor tissues from patients previously recruited in Qidong County, China, with approval by the IARC Ethics Committee (project ref. no. 15-20) (Supplementary table 11). All 6 HCCs harbored the *TP53* R249S somatic mutation, and the patients were positive for HBsAg and negative for hepatitis C infection (Szymanska et al. 2009).

### Sources of previously reported somatic mutation data from human tumors

We obtained previously reported sequencing read data as follows: WGS reads from (Sung et al. 2012), ftp.ncbi.nlm.nih.gov:/sra/sra-instant/reads/ByStudy/sra/ERP/ERP001/ERP001196/; WES reads from the recent African immigrants in (Schulze et al. 2015), European Genome-Phenome Archive (EGAS00001001002); TCGA WES reads reported in (Schulze et al. 2015) from the (National Cancer Institute Genomic Data Commons Data Portal); somatic variant calls from the additional TCGA (The Cancer Genome Atlas 2017) WES sequences from sftp://tcgaftps.nci.nih.gov/tcgajamboree/mc3/pancan.merged.v0.2.7.PUBLIC.maf.gz downloaded on Nov 11, 2016. TCGA WGS reads from DO23048 from CGHub (https://cghub.ucsc.edu/). For analysis of evidence of AFB1 exposure among TCGA HCCs treated in North America, we obtained clinical information from https://portal.gdc.cancer.gov on Apr 15, 2017. We considered HCCs that were definitely treated in North America. To this end, we excluded HCCs obtained from biorepositories from numerator and denominator, as we were unable to determine where they were treated. In particular we excluded the 4 TCGA HCCs reported in (Schulze et al. 2015) as "Asian" (CCA5UE, CCA7IG, CCA7II, CCA7IL), as these were obtained from a biorepository. We obtained previously reported somatic variant calls from (Fujimoto et al. 2016) from the International Cancer Genome Consortium data portal (https://dcc.icgc.org/).

### Whole-genome sequencing of human cell lines and mouse tumors

For DNA extraction from frozen mouse tissue, we placed 15 to 25mg of frozen tissue in a 2ml microcentrifuge tube containing ATL buffer (220 μl, Qiagen) and Proteinase K (20 μl, Qiagen). Tissue was homogenized in the TissueLyser II (speed:20-30Hz, Qiagen) for 1-2 minutes then placed into thermomixer (56 ^o^C, shaking speed: 900rpm, Eppendorf) for 3 hours. After homogenization, 220 μL of supernatant was transferred to another sample tube containing RNase A (4 μl, Qiagen) and incubated at room temperature for 2 minutes before DNA extraction using QIAsymphony (Qiagen). For DNA extraction from cell lines, cells were trypsinized, washed with PBS and centrifuged to form a pellet. We then added a mixture of buffer ATL and Proteinase K and extracted DNA using QIAsymphony (Qiagen). For both tissue and cell-line DNA, quality was assessed by an Agilent 2200 T before sequencing. Supplementary table 12 provides details on the sequencing runs.

### Whole-exome sequencing of human FFPE samples

Genomic DNA was extracted from macro-dissected paraffin sections, and 250 nanograms of each DNA sample were sheared by Covaris (Covaris, Inc.) to ∼300 bp fragments, as previously described (Castells et al. 2015). We prepared libraries with the Kapa LTP Library Preparation Kit (Kapa Biosystems) according to the manufacturer’s recommendations. Four libraries (250 ng each) were pooled per capture with the Nimblegen Roche SeqCap EZ Exome v3 reagent (Roche), and the exome-enriched mix was PCR-amplified in 10 cycles. The post-enrichment material was diluted in 420 mL water to a final concentration of 6 pmol/L and sequenced as described in Supplementary table 7.

### Alignment and variant calling

For whole genome sequencing (WGS) we used BWA mem (v0.7.12) (Li and Durbin 2009) with the –R and –M options to align reads to hs37d5 (human) or mm10 (mouse), followed by sorting, PCR duplicate removal and merging using Sambamba (v0.5.8) (Tarasov et al. 2015). Strelka (v1.014) (Saunders et al. 2012) called somatic variants, with non-default parameters as follows: ssnvNoise=0.00000005, minTierMapq=15, ssnvQuality_LowerBound=25, sindelQuality_LowerBound=20 and isWriteRealignedBam=1; any variants with variant depth / total depth < 0.1 were removed. Within each cell line (HepaRG and HepG2) we used Strelka to call "somatic" variants in each clone, *c*, against the other two clones, and considered the intersection of the "somatic" variants as the mutations specific to *c*. For tumors from mice M1, M2, and M3, which were treated with AFB1 only, Strelka called variants from reads from the tumor and matched non-malignant tissue. For mice M4, M5, and M6, which carried the HBsAg transgene and were treated with AFB1, we used Strelka to call "somatic" variants in each tumor, *t*, against the other two tumors, and considered the intersection of the "somatic" variants as the somatic mutations in *t.*

For WES data, BWA mem (v0.7.9a) with default parameters aligned reads to hs37d5, followed by samtools (v 0.1.8 r613) (Li et al. 2009) to sort and reads and remove duplicates. GATK (v2.2-25-g2a68eab) was used for local realignment around indels and base quality recalibration (McKenna et al. 2010). Picard tools corrected mate pair information, and MuTect (v 1.1.4) (Cibulskis et al. 2013) with default parameters called variants on tumor and matched normal pairs. We restricted analysis to mutations in the capture target.

### Additional filtering of mutations

For human samples, variants in dbSNPv132, 1000 genomes (1000 Genomes Project Consortium 2015), segmental duplications, microsatellites and homopolymers, and the GL and decoy sequences were excluded. For mouse tumors, variants in dbSNPv142, regions of segmental duplication, or regions of tandem repeats were excluded. For human and mouse WGS data, we also excluded candidate somatic variants at sites where ≥ 2 normal samples each contained ≥ 2 reads with a variant base.

### Expression data for analysis of transcription strand bias

We obtained expression data as follows: HepaRG, (average of accessions GSM1139508, GSM1139509, GSM1139510 from Gene Expression Omnibus, www.ncbi.nlm.nih.gov/geo/) (Ambolet-Camoit et al. 2015); HepG2, Epigenomic Roadmap (http://egg2.wustl.edu/roadmap/data/byDataType/rna/expression/57epigenomes.RPKM.pc.gz); normal human liver, Epigenomic Roadmap; normal mouse liver, (http://chromosome.sdsc.edu/mouse/download/19-tissues-expr.zip)(Shen et al. 2012).

### Analysis of replication timing and replication strand bias

For analysis of human cell lines and primary tumors, we obtained processed replication timing (RepliSeq) data that provided early and late replication regions for HepG2 from Gene Expression Omnibus, (www.ncbi.nlm.nih.gov/geo/, accession GSM923446). We determined replication strand based on the local maxima and minima of wavelet-smoothed signal data for HepG2 (Liu et al. 2015); we took the peaks as replication initiation zones and the valleys as replication termination zones. We inferred replication direction as proceeding from initiation to termination zones.

### Principal components analysis

We used the R function “prcomp” for principal components analysis (PCA) (R Development Core Team 2017). Trinucleotide frequencies in the human genome, human exome and mouse genome are different. Therefore, to compare mutation spectra from these three sources, the spectra of human exomes and mouse genomes were normalized to human genome frequencies, as is a common practice in this field.

### Analysis of variant allele frequencies in the context of tumor purity and aneuploidy

Tumor purity (proportion of malignant cells in the tumor), ploidy estimation, and copy number and depth ratios across each mouse tumor genome were calculated and visualized using Sequenza (Favero et al. 2015). In brief, we created Sequenza input files by applying bam2seqz to the the tumor and normal BAMs file along with information on the, reference genome. We generated binned input with seqz-binning with -w 500, and then it loaded into R using sequenza.extract() function from the R sequenza library. Tumor content, ploidy, and copy number and depth ratios were calculated using sequenza.fit(). We plotted results with sequenza.results(). Custom scripts were used to separate SNVs of each mouse tumor according to mutation category, VAF, and locus copy number. Probability density histograms at were created using the R function hist() with probability=T; probability density lines were estimated using the R density() function (R Development Core Team 2017).

### In-silico decomposition of aflatoxin-related mutational signatures and assignment of mutational-signature activities to HCCs

To discover AFB1-related mutational signatures in experimental data and somatic mutation data from human HCCs selected for likely aflatoxin exposure, we used the nsNMF method from R NMF package (https://cran.r-project.org/web/packages/NMF/index.html) with the sparsity parameter theta = 0 and nrum=200. We studied G > N signatures in trinucleotide context in the experimental AFB1 data and in HCCs likely to have been exposed to aflatoxin. Mouse spectra were normalised the trinucleotide frequencies of the human genome. In addition, because of the possible presence of a signature similar to COSMIC Signature 5 in the low-VAF somatic mutations in some of the mouse tumors (Supplementary figures 4-9), for the AFB1-only mice we restricted this analysis to mutations with VAFs > 0.2 to enrich for mutations due only to AFB1. We studied G > N signatures (rather all 96 mutations in trinucleotide context) to avoid interference from other mutational processes that affect A > N mutations, including COSMIC Signatures 12, 16, and 17 (Wellcome Trust Sanger Institute 2016).

To assess evidence of AFB1 exposure in human HCCs, we developed a customized method based on the Lee and Seung NMF algorithm(Lee and Seung 1999) in which we fixed the W (signature) matrix and only updated the H (exposure) matrix, while incorporating the smoothing matrix from (Pascual-Montano et al. 2006). We used this method with the R NMF package, and the code is available on request. We applied this to human HCCs with the set of signatures previously found in HCCs (COSMIC Signatures 1, 4, 5, 6, 12, 16, 17, 22, 23 (Wellcome Trust Sanger Institute 2016)), plus the two AFB1-related signatures identified in this study, with sparsity parameter theta = 0.

### *P* values for generation of a mutational spectrum from a signature

We calculated *p* values for the observation of a particular observed mutational spectrum, *s*, consisting of *n* mutations, under the null hypothesis that it was generated from a particular mutational signature as follows. We used the R rmultinom function create 10,000 synthetic spectra of *n* mutations each, drawn with the probabilities specified by the mutational signature. For each replicate we calculated the cosine similarity between the synthetic spectrum and the signature. We took as the *p* value the proportion of replicates in which the cosine similarity was less than the cosine similarity between the signature and the observed spectrum, *s*.

## List of abbreviations

AFB1: aflatoxin B1
AFB1sig: Extension of AFB1sig_G>N_ to all 96 trinucleotide contexts by setting A > N mutations to 0
AFB1sig_G>N_: 1st NMF-extracted signature from G>N mutations in trinucleotide context experimental data and likely strongly aflatoxin exposed HCCs
AFsig2: Extension of AFsig2_G>N_ to all 96 trinucleotide contexts by setting A > N mutations to 0
AFsig2_G>N_: 2nd NMF-extracted signature from G>N mutations in trinucleotide context experimental data and likely strongly aflatoxin exposed HCCs
HbsAg: hepatitis B surface antigen
HCC: hepatocellular carcinoma
HepG2: a human hepatoblastoma-derived cell line used in this study
HepaRG: a human HCC-derived cell line that was differentiated to cells resembling hepatocytes
IARC: International Agency for Research on Cancer
IC50: half maximal inhibitory concentration
NMF: nonnegative matrix factorization
PC1: principal component 1
PCA: principal components analysis
TCGA: The Cancer Genome Atlas project (https://cancergenome.nih.gov/)
WES: whole-exome sequencing
WGS: whole-genome sequencing

## Authors’ contributions

MNH, WY, AB, KS, JZ, SGR drafted the manuscript and prepared figures. WWT under the supervision of KS generated the mouse models and performed all in vivo experiments. AJ, SSM, RO, SP carried out other laboratory studies. MNH, WY, MA, AN, carried out electronic analyses. BA-A, SV, AH analyzed and organized human tumor samples. MNH, WY, AB, MO, MH, PT, BTT, KS, JZ, SGR designed the study and edited the manuscript.

## Acknowledgements

The results published here are in part based upon data generated by the TCGA Research Network: http://cancergenome.nih.gov.

## Data availability

Whole genome sequencing reads for the human cell lines, mouse tumors, are available at the European Genome-Phenome Archive (https://www.ebi.ac.uk/ega/home) under accession number EGAS00001002162. Due to IRB protocol restrictions, whole exome reads from HCCs and non-tumor samples from Qidong are available upon request made to J.Z.

